# Examining the Thermotropic properties of Large, Circularized Nanodiscs

**DOI:** 10.1101/2025.04.07.647641

**Authors:** Mark J. Arcario, Vikram Dalal, David Fan, Wayland W.L. Cheng

## Abstract

Nanodiscs, soluble membrane mimetics composed of an amphipathic membrane scaffold protein encircling a lipid bilayer, are widely used in biophysical and structural studies of membrane proteins. Because many membrane proteins are responsive to their membrane environment, through specific protein-lipid interactions and bulk membrane shape and structure, it is important to understand the properties of lipid bilayers contained within nanodiscs in order to interpret studies using this technology. Nanodiscs are known to alter lipid properties, such as membrane thickness and melting temperature, and interactions with the nanodisc rim have been hypothesized to produce local perturbations in lipid structure and dynamics. Larger nanodiscs should compensate for this effect with a larger unperturbed area. To test this hypothesis, we examined the lipid bilayer properties of several lipids (DMPC, DPPC, POPC, DSPC) and soy polar extract in circularized nanodiscs of 11 nm to 50 nm diameter using the environmentally-sensitive fluorophore, Laurdan. In nanodiscs containing a single lipid type, as nanodisc size increased, lipid packing, melting temperature, and cooperativity better approximated the properties of that lipid in large unilamellar vesicles (LUVs). In spNW50 (50 nm nanodisc), the lipid packing and melting temperature were identical to LUVs. However, nanodiscs containing soy polar lipids did not follow this trend suggesting that complex lipid mixtures may produce preferential incorporation of lipids into the nanodisc or nonhomogeneous distribution of lipids within the nanodisc.

## 1. Introduction

Membrane proteins are challenging targets for biophysical and structural studies because of the difficulty of stabilizing the hydrophobic transmembrane domain throughout purification and analysis. Detergents or amphipols can be used to solubilize membrane proteins, but the process of solubilization removes surrounding lipids. In many cases, these lipids are necessary for the structural integrity of the protein or regulate protein function by direct binding or altering membrane properties [1]. In some cases, detergent-solubilized membrane proteins have been shown to have altered structure and function [2, 3]. Nanodiscs, discoidal bilayers stabilized by an amphipathic scaffold, are a useful alternative membrane mimetic that maintains a lipid bilayer structure while still allowing access to both the intracellular and extracellular domains of membrane proteins.

The original nanodiscs consist of a lipid bilayer encircled by membrane scaffolding proteins (MSPs), an engineered derivative of human apolipoprotein A-1 [4]. This concept has since been applied other proteins [5, 6], DNA [7], and copolymeric [8–10] scaffolds. Nanodiscs have become the preferred platform for use in structural studies [11], such as cryo-electron microscopy (cryo-EM) [12] and nuclear magnetic resonance (NMR) [13], which has fueled a significant increase in the number of available membrane protein structures. Functionalization of nanodiscs, including “dark” nanodiscs [14], affinity tags for surface immobilization [15], anti-nanodisc antibodies [16] and addition of Förster resonance energy transfer pairs [17, 18] has made them a useful platform in a wide range of biophysical techniques.

While nanodiscs stabilize a soluble lipid bilayer, alterations in the physicochemical properties of the lipid bilayer have been demonstrated, including melting temperature [19, 20], bilayer height [19, 21–24], membrane bending modulus [23], area per lipid [21–23, 25], lipid chain order [20–22, 25–27], configurational entropy [25], and two-dimensional diffusion [21, 25]. Perturbations to these lipid properties have challenged the utility of MSP-based nanodiscs. While copolymer-based nanodiscs can directly extract membrane proteins and lipids from native membranes, similar differences in lipid structure and dynamics have been observed in copolymeric nanodiscs [28, 29].

The differences in lipid physicochemical properties have been attributed to a complex interplay of nanodisc geometry, nanodisc-lipid stoichiometry, and lateral forces induced by the thermal expansion-contraction of the nanodisc scaffold.

Computational [21–23, 26] and structural [26] studies of nanodiscs report that the physical properties of the lipids are spatially heterogenous, driven by the geometry and dynamics of lipids at the nanodisc rim (*i*.*e*., where the lipid bilayer meets the MSP construct). Denisov *et al*. [19], attributed the decreased enthalpy change of the gel-liquid phase transition in nanodiscs to perturbed boundary lipids close to the nanodisc rim. Comparing the enthalpy change of the gel-liquid transition in nanodiscs versus liposomes, it was suggested that the width of the perturbed boundary layer was a fixed ∼ 1.5 nm (approximately two radial layers of lipid at the nanodisc rim) and that larger nanodiscs would have properties indistinguishable from an unperturbed lipid bilayer. This calculation employed a model which assumed lipids existed either as “perturbed” or “unperturbed”, but some studies [26, 27] have suggested that lipids in nanodiscs exist on a continuum of perturbation. Additionally, simulations comparing smaller (MSP1, ∼9.8 nm diameter) and larger (MSP2N2, ∼18.4 nm diameter) nanodiscs showed that the geometry induced by the nanodisc rim propogates further into the bilayer for the larger nanodisc, although the larger nanodisc was simulated for only 150 ns [23]. Therefore, it remains unclear if larger nanodiscs are able to reproduce properties of unperturbed lipid bilayers.

Until recently, nanodisc scaffolds were stable only to a diameter of approximately 15-18 nm [30]. Engineering of the scaffold to contain sites for covalent circularization, via the sortase-mediated method [31], split intein method [32], or SpyCatcher-SpyTag technology [33], has produced thermally-stable nanodiscs up to 100 nm. The wide range of nanodisc sizes now available (6-100 nm) makes it possible to test whether lipid properties in larger nanodiscs better approximate an unperturbed lipid bilayer. Here, we use the solvatochromatic, lipophilic fluorophore, Laurdan (Fig. 1), to measure lipid packing in nanodiscs ranging from 11-50 nm and compare this to lipid packing in large unilamellar vesicles (LUVs) of the same lipid composition, focusing on four lipids: DMPC (1,2-dimyristoyl-*sn*-glycero-3-phosphatidylcholine), DPPC (1,2-dipalmitoyl-*sn*-glycero-3-phosphatidylcholine), POPC (1-palmitoyl-2-oleoyl-*sn*-glycero-3-phosphatidylcholine), and DSPC (1,2-distearoyl-*sn*-glycero-3-phosphatidylcholine). We observe that as nanodisc size increases, the properties of the lipid bilayer better approximate those of LUVs, with spNW50 (50 nm nanodisc) appearing identical to LUVs in the case of DPPC, POPC, and DSPC. For all saturated lipids examined, melting temperature in the spNW50 nanodisc was identical to that of LUVs and cooperativity of the phase transition increased as nanodisc size increased. Overall, our results demonstrate that lipid bilayers in larger nanodiscs more closely resemble an unperturbed lipid bilayer in LUVs. The effects of nanodisc size on lipid bilayer properties are likely to impact the structure and function of some membrane proteins when reconstituted in nanodiscs.

**Figure 1:**
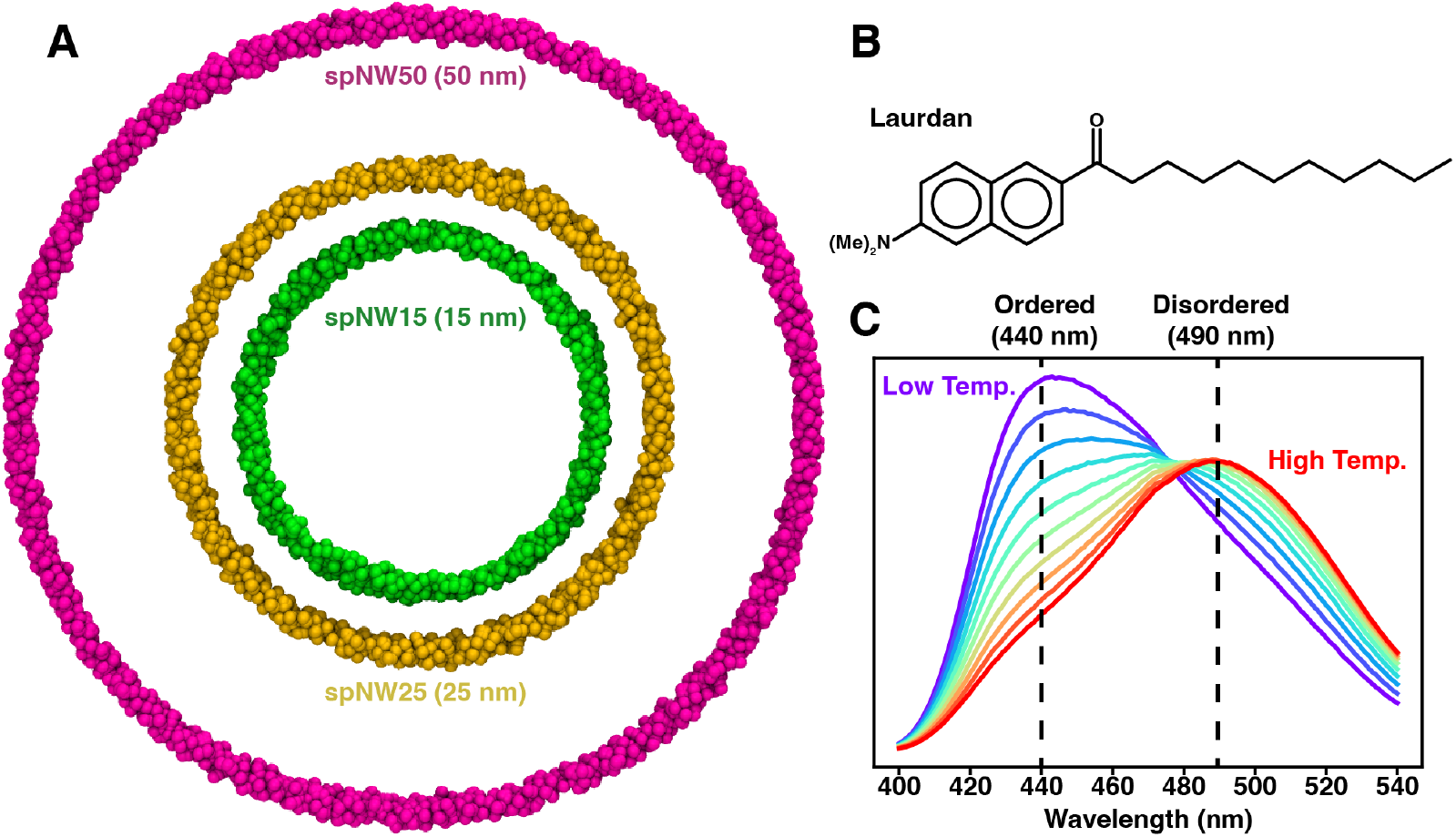
(A) Molecular models of the nanodisc MSP scaffold showing the relative size of each nanodisc with the scaffold shown as van der Waals spheres (spNW15, dark green; spNW25, gold; spNW50, magenta). spMSP1D1 is omitted in this figure as the scaffold would overlap spNW15 significantly. The chemical structure of Laurdan is shown (B) along with a representative set of Laurdan emission spectra (C) from low temperature (purple, below the transition temperature of the membrane) to high temperature (red, above the transition temperature of the membrane). The peak emission is 440 nm when lipids are in an ordered phase and 490 nm when lipids are in a disordered phase.

## 2. Materials and Methods

### 2.1. Materials

1,2-dimyristoyl-*sn*-glycero-3-phosphatidylcholine (DMPC), 1,2-dipalmitoyl-*sn*-glycero-3-phosphatidylcholine (DPPC), 1-palmitoyl-2-oleoyl-*sn*-glycero-3-phosphatidylcholine (POPC), 1,2-distearoyl-*sn*-glycero-3-phosphatidylcholine (DSPC), and 1-(6-(dimethylamino)naphthalen-2-yl)dodecan-1-one (Laurdan) were obtained from Avanti Polar Lipids (Alabaster, AL). Plasmids for spMSP1D1 (AddGene ID 173482), spNW15 (AddGene ID 173483), spNW25 (AddGene ID 173484), and spNW50 (AddGene ID 173486) were purchased from Addgene (Watertown, MA). BioBeads SM-2 adsorbent beads were purchased from Bio-Rad Laboratories (Hercules, CA). The quartz cuvette used for fluorometric assay was purchased from Starna Cells (Atascadero, CA). All other chemicals were purchased from Sigma-Aldrich (St. Louis, MO).

### 2.2. Production of large unilamellar vesicles

Laurdan powder was dissolved in a 1:1 solution of chloroform and methanol to a concentration of 1 mM and stored at -20°C until use. Using a glass syringe at room temperature, Laurdan was added in a 1:200 ratio to the stock solution of lipid in a glass vial. The resulting solution was dried under a gentle stream of N_2_ and placed in a vacuum desiccator overnight to remove any residual solvent, leaving a thin lipid film. This lipid film was then hydrated with buffer A (20 mM HEPES, 150 mM NaCl, pH 7.0) at room temperature and vortexed for 5 minutes. Each sample then underwent five freeze-thaw cycles. Samples were flash frozen in liquid nitrogen for 3 minutes and thawed by placing in an aluminum heating block. During the thawing process, the aluminum heating block was maintained above the lipid’s melting temperature (room temperature for POPC, DMPC, and soy polar extract, 50°C for DPPC, and 70°C for DSPC). Following five freeze-thaw cycles, the suspension was extruded twenty times at room temperature through a polycarbonate membrane with a nominal pore size of 0.1 *µ*m (Avanti Polar Lipids, Alabaster, AL). Large unilamellar vesicles (LUVs) were stored at 4°C until use.

### 2.3. Expression and Purification of Circularized Nanodiscs

Expression and purification of the nanodiscs followed the published protocol [33]. spMSP1D1, spNW15, spNW25, and spNW50 plasmids were transformed into *E. coli* BL21STAR (DE3) cells and grown in LB media at 37°C with kanamycin (50 *µ*g/mL) until an optical density of ∼ 0.7 at 600 nm. Protein expression was induced with 0.2 mM IPTG and the cultures maintained at 16°C overnight. Cells were pelleted by centrifugation at 3450g for 20 minutes at 4°C. The supernatant was removed and the cells were stored at -80°C until purification. For purification, the cell pellet was resuspended in buffer B (50 mM Tris-HCl at pH 8, 100 mM NaCl, 5% glycerol, 2 mM *β*-mercaptoethanol) and lysed using the EmulsiFlex system (15,000 psi for 3 passes). Cell lysate and other insoluble material was separated by centrifugation at 12,000g for 45 minutes at 4°C. The supernatant was loaded onto a Ni-NTA column (pre-equilibrated with ten column volumes of buffer B) and washed with twenty column volumes of 50 mM Tris-HCl at pH 8, 20 mM Imidazole, 400 mM NaCl, 5% glycerol, 2mM *β*-mercaptoethanol. The protein was eluted with 50 mM Tris-HCl at pH 8, 500 mM Imidazole, 400 mM NaCl, 5% glycerol, 2 mM *β*-mercaptoethanol and stored at -80°C.

### 2.4. Formation of Circularized Nanodiscs

Laurdan (as prepared in 2.2) was added in a 1:200 ratio to the stock solution of lipid in a glass vial at room temperature. The resulting solution was dried under a gentle stream of N_2_ and placed in the vacuum desiccator overnight to remove any residual solvent, producing a thin lipid film. Buffer A with 50 mM cholate at room temperature was added to the glass vial containing the thin lipid film to a lipid concentration of 20 mM. The solution was vortexed at room temperature for 5 minutes, providing complete dissolution of the lipid film for POPC and DMPC. In the case of DPPC and DSPC, the solution containing lipid in buffer A with 50 mM cholate was heated above the melting temperature (as in 2.2) and sonicated for 10 minutes following the initial vortexing to completely solubilize the lipids. The resulting solution was allowed to equilibrate at room temperature. Appropriate volumes of circularized nanodisc scaffold protein (spMSP1D1, spNW15, spNW25, or spNW50) and solubilized lipids (molar ratios of 1:90 for spMSP1D1, 1:150 for spNW15, 1:350 for spNW25, and 1:1000 for spNW50) were combined and rotated for 1 hour above the melting temperature of the lipid (4°C for POPC, 30°C for DMPC, 50°C for DPPC, and 70°C for DSPC). BioBeads were added to the solution and rotated at the same temperature overnight (rotated 3 hours for DPPC and DSPC given the elevated temperature) in order to remove cholate and allow self-assembly of the nanodisc. Following this, the solution was removed from the BioBeads and centrifuged at 20,000 rpm for 10 minutes at 4°C. The supernatant was withdrawn into a glass syringe and purified by size exclusion chromatography on an Äkta Pure System (Cytiva Life Sciences, Marlborough, MA) using a Superose 6 Increase 10/300 column (Cytiva Life Sciences) pre-washed and equilibrated with Buffer A at 4°C. The fraction containing the elution peak (Fig. 2, Table 1) was stored at 4°C until use.

**Table 1:**
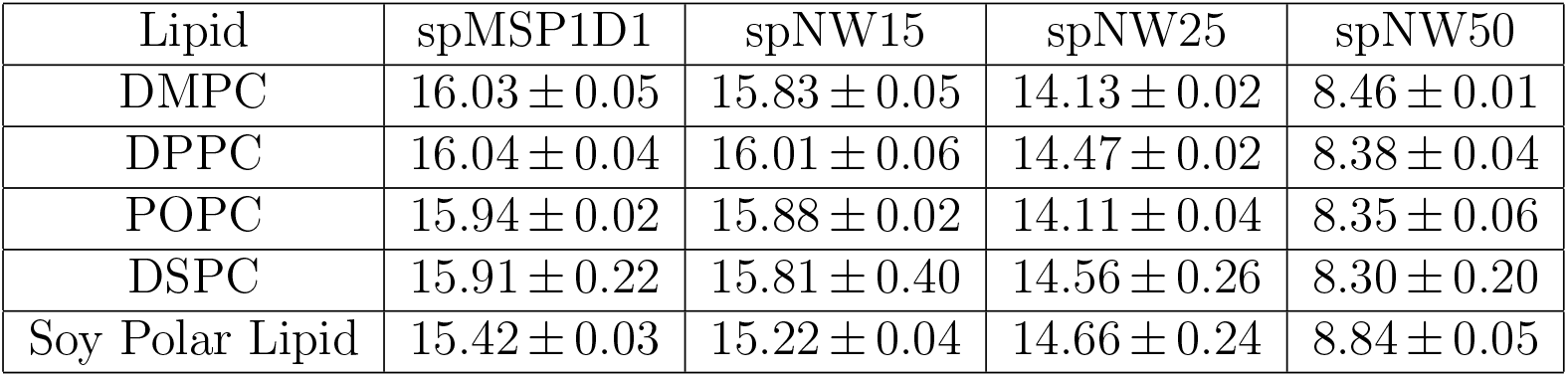
Elution volumes for lipid-filled nanodiscs. This table shows the elution volume (in mL) for each nanodisc size and lipid condition. Values presented are average (*±* standard deviation) across the three samples used in the analysis.

**Figure 2:**
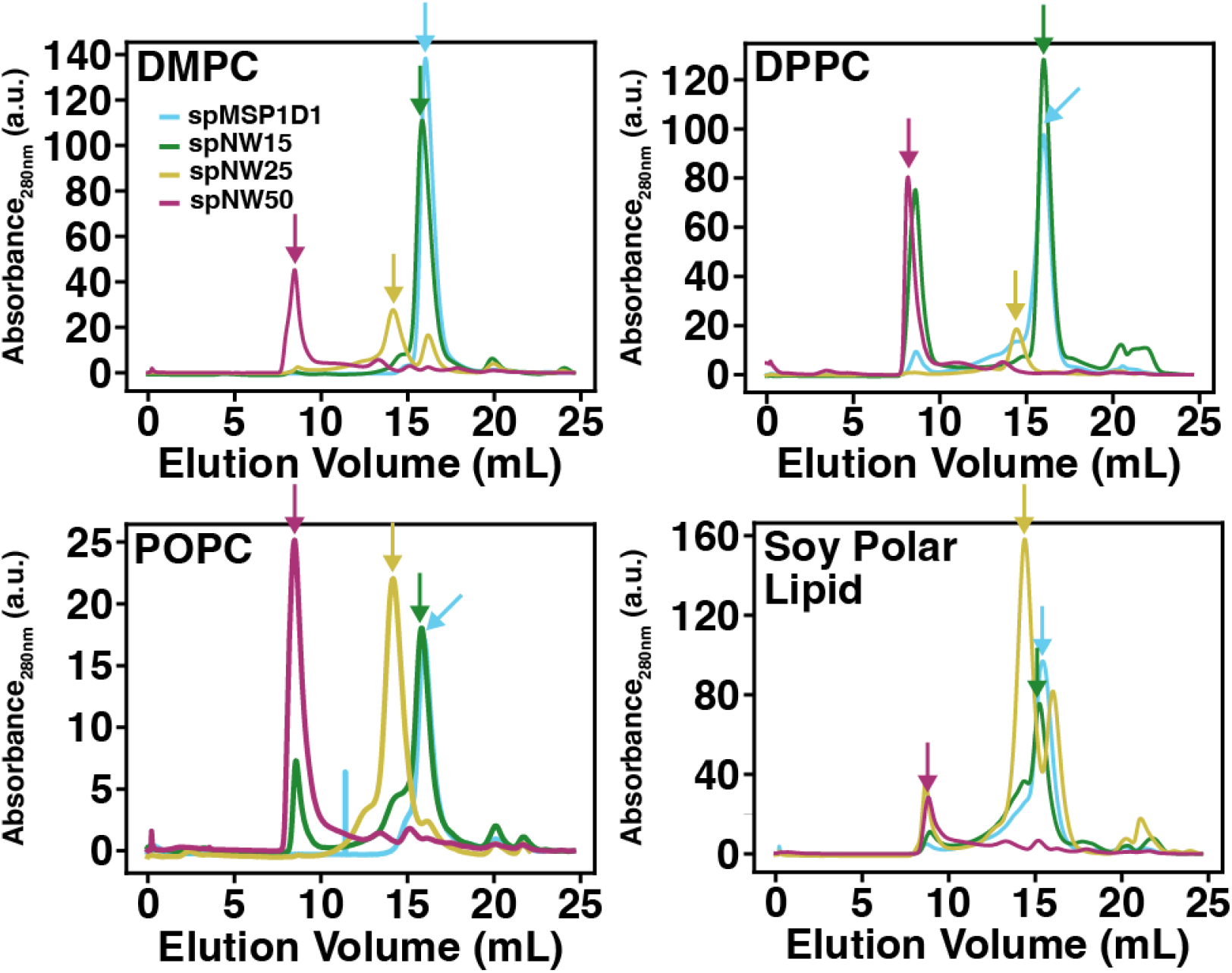
Size exclusion chromatography for lipid-filled nanodiscs containing POPC (*left*), DMPC (*middle left*), DPPC (*middle right*), and soy polar lipid (*right*). For each plot, the absorbance of the sample at 280 nm is shown as a function of elution volume (mL) from the column for each nanodisc sample (spMSP1D1, light blue; spNW15, dark green; spNW25, gold; spNW50, magenta). Arrows denote the peaks collected for fluorometric assay.

### 2.5. Fluorescence Spectroscopy Measurements

Steady state emission spectra for each sample were acquired using a Fluoromax+ spectrophotometer (Jobin Yvon, Edison, NJ) with temperature control achieved via Peltier thermocouple (Model TC-1 from Quantum Northwest, Lady Lake, WA). Samples were placed in quartz cuvettes and incubated at the starting temperature for 5 minutes. For samples in which the temperature was raised by more than 1°C between measurements (*i.e*., POPC and soy polar extract), the sample was incubated at each new temperature for 5 minutes before the subsequent measurement. For samples in which the temperature was raised by 1°C between measurements (*i.e*., DMPC, DPPC, and DSPC), the sample was incubated for 3 minutes at each new temperature before the subsequent measurement. Laurdan was excited at 385 nm with a bandwidth of 1.2 nm on both the excitation and emission monochromators. Steady state spectra were acquired in 1 nm steps from 400 nm to 540 nm using an integration time of 0.2 s. Within an individual sample, each steady state spectrum represents a simple average of three scans using the above parameters. Fluorescence spectra were analyzed by calculating Laurdan generalized polarization (GP) according to:

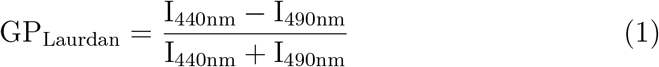

where I_440nm_ is the intensity of the steady state spectrum at 440 nm and I_490nm_ is the intensity of the steady state spectrum at 490 nm. Data was visualized using an in-house Python (version 3.6.8) script available at https://github.com/mjarcario/Laurdan-Data-Extraction/tree/main. The derivative of the temperature-dependent GP plots was calculated using the central finite difference method, available in the same Python script.

## 3. Results

To test the idea that larger nanodiscs are less perturbing to the lipid bilayer, we used the environmentally-sensitive membrane dye, Laurdan, to report on lipid packing in nanodiscs ranging from 11 nm to 50 nm in diameter (Fig. 1) and compared this with large unilamellar vesicles (LUVs) of the same lipid composition. The chosen range of nanodisc sizes span from those originally described by Sligar and colleagues (11 nm) [4] to sizes only recently made possible via covalent circularization of the nanodisc scaffold (50 nm) [33]. Laurdan has been extensively characterized [34] and routinely used as a reporter of the physical properties of lipids contained in nanodiscs [19, 29, 35–37] (in addition to its well-documented use in liposomes) due to its sensitivity to the polarity of the local environment [38]. Specifically, the emission peak of the dye undergoes a red shift from 400 nm to 490 nm (Fig. 1) response to increased aqueous solvation [34], which is associated with decreased lipid packing and decreased membrane order [38]. The normalized relative intensity of these two peaks, called the generalized polarization (GP, see 2.5 for the definition), quantifies the degree of membrane packing with higher values indicating tighter lipid packing and lower values indicating looser lipid packing.

In this study, we chose four lipids whose properties are well-characterized (see [39] for a thorough review) and had been previously examined in smaller nanodiscs [19, 29, 36], namely DMPC (di-14:0), DPPC (di-16:0), POPC (16:0,18:1), and DSPC (di-18:0). In addition, we investigated the properties of soy polar lipid (asolectin), which is widely used in structural studies by cryo-EM [40–45]. To ensure that all measurements contain only well-formed nanodiscs, we isolated each nanodisc sample using size exclusion chromatography. For all lipid types, each nanodisc preparation produced monodisperse peaks (Fig. 2) at elution volumes consistent with those previously published [33]. All fluorescence measurements were collected from a single fraction containing only the peak of interest.

### 3.1. spNW50 recovers bulk lipid properties of large unilamellar vesicles for single lipid compositions

The saturated lipids studied here (DMPC, DPPC, and DSPC) have transition temperatures at 24°C, 41°C, and 54°C respectively, which allows investigation of lipid packing in both ordered and disordered phases (Fig. 3). The LUVs of DMPC, DPPC, and DSPC showed a reproducible, sharp phase transition at the expected transition temperature (Supplemental Figs. S1-S3).

**Figure 3:**
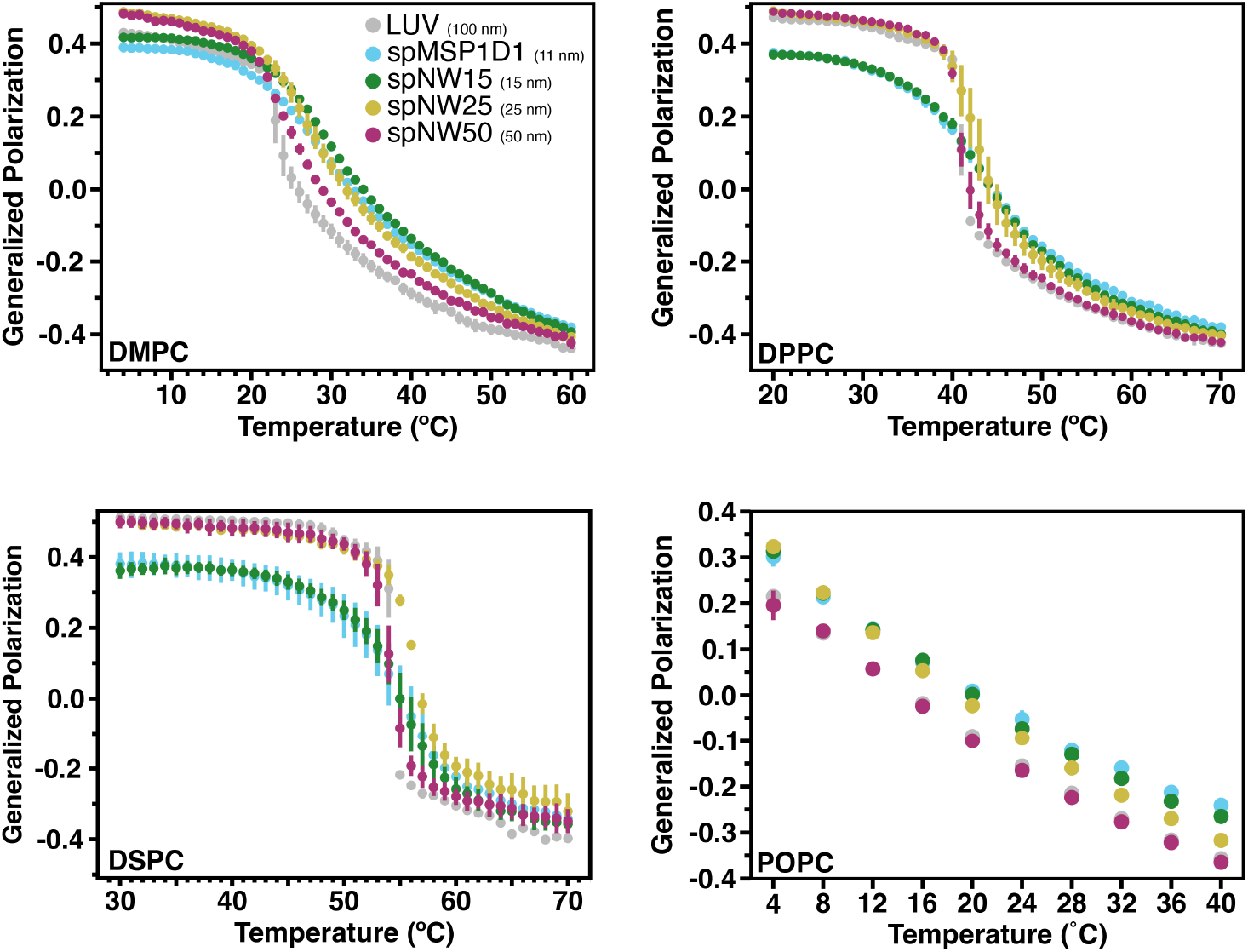
Generalized polarization (GP) values as a function of temperature for model lipid types DMPC (*top left*), DPPC (*top right*), DSPC (*bottom left*) and POPC (*bottom right*). Higher GP values signify tighter packing of the membrane, whereas lower GP values signify looser packing. Each data point represents the average of three independent measurements with the error bars representing the standard deviation of the the three measurements. Data for nanodiscs are colored as light blue for spMSP1D1 (11 nm), dark green for spNW15 (15 nm), gold for spNW25 (25 nm), and magenta for spNW50 (50 nm). Data for LUVs are colored gray. Where data for spMSP1D1 (light blue) is not visible in these plots, data from spNW15 (dark green) overlie due to similarity in the GP values and size of data points.

For DMPC, the two smaller diameter nanodiscs, spMSP1D1 (11 nm) and spNW15 (15 nm), demonstrate lipid packing similar to LUVs at temperatures below the phase transition, but tighter lipid packing (*i.e*., higher GP values) compared to LUVs above the phase transition. In contrast, DMPC in larger nanodiscs, spNW25 (25 nm) and spNW50 (50 nm), demonstrate tighter lipid packing compared to LUVs below the phase transition, but lipid packing similar to LUVs above the phase transition. All DMPC-filled nanodiscs, however, have a less cooperative phase transition evidenced by the shallow decrease in GP values as a function of temperature (Fig. 3). This “continuous phase transition” has been demonstrated previously in both MSP-based and polymeric nanodiscs [19, 29, 35, 46] and attributed to perturbations induced by the nanodisc rim. As nanodisc size increases, the transition becomes sharper, indicating a more cooperative transition in larger nanodiscs compared to smaller nanodiscs.

The derivative of the temperature-dependent GP plot can yield information about the phase transition, including transition temperature (location of the peak), relative cooperativity of the phase transition (width at half-maximum amplitude of the peak), and enthalpy of the phase transition (area under the curve). This approach has been used previously with multilamellar vesicles of single lipids and has demonstrated good agreement with differential scanning calorimetry measurements [47]. For DMPC, nanodiscs with smaller diameter show an increased melting (main transition) temperature (T_*M*_) compared to LUVs (Fig. 4), which has been demonstrated previously in non-circularized nanodiscs [19, 35]. In contrast, the phase transition temperature decreased in polymer-based nanodiscs containing saturated phosphatidylcholine lipids [46]. Interestingly, as the size of the nanodisc increases, T_*M*_ decreases and at 50 nm coincides with that of LUVs of the same composition (Fig. 4). However, the phase transition is much broader for all nanodiscs compared to LUVs, indicating a less cooperative phase transition in nanodiscs compared to LUVs, consistent with the fact that the nanodisc rim perturbs local lipid-lipid interactions. Again, as nanodisc diameter increases, the cooperativity of the phase transition increases, but even at 50 nm, cooperativity is lower than in LUVs. Overall, similar trends are noted for DPPC- and DSPC-containing nanodiscs in regards to T_*M*_ (Fig. 3) and cooperativity (Fig. 4). As noted in DMPC-containing assemblies, lipids in nanodiscs with smaller diameters have a right-shifted main transition temperature as compared to an LUV with the same lipid composition (Fig. 4). As nanodisc size increases, the transition temperature shifts leftward and the cooperativity of the phase transition increases. The transition temperature for DPPC, however, is less affected overall by incorporation into nanodiscs than DMPC (Fig. 5), with the ΔT_M_ between LUV and spMSP1D1 being 2°C in DPPC-containing nanodiscs compared to 5°C in DMPC-containing nanodiscs; this observation was also made in small, non-circularized nanodiscs previously [19]. Further, DSPC shows less deviation in the main transition temperature than DPPC (Fig. 5), with even the smallest nanodisc (spMSP1D1) showing the same main transition temperature as a DSPC LUV. As in DMPC, when compared to smaller nanodics, larger nanodiscs are more ordered at temperatures below the transition point and less ordered at temperatures above the transition point. In contrast to DMPC, larger nanodiscs containing DPPC or DSPC better approximate LUVs at temperatures both above and below the transition point and, in the case of spNW50, display the same order across the entire temperature spectrum. Taken together, these results suggest that lipids incorporated into the nanodisc may be accommodated by the scaffold differently depending on the length and geometry of the lipid acyl chains.

**Figure 4:**
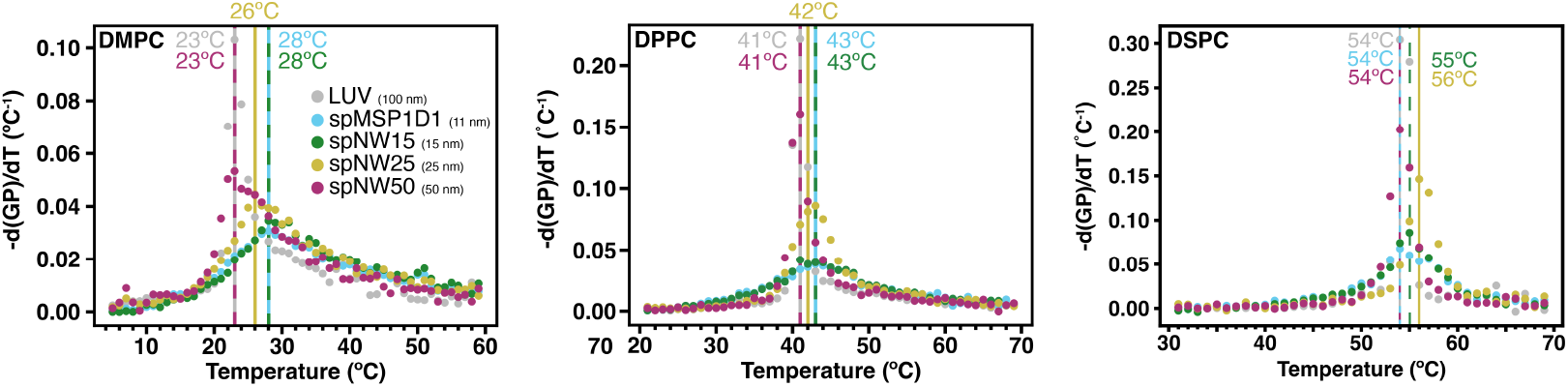
Transition temperatures of DMPC (*left*), DPPC (*middle*), and DSPC (*right*) as measured by the negative derivative of generalized polarization (GP) values with respect to temperature using the central difference algorithm applied to the average of three independent experiments. Here, the temperature at which the derivative is maximum represents the transition temperature, which is marked with a vertical line with the same color as the data points (spMSP1D1, light blue; spNW15, dark green; spNW25, gold; spNW50, magenta; LUV, gray). The transition temperature for each nanodisc and LUV is also shown with a number of the same color.

**Figure 5:**
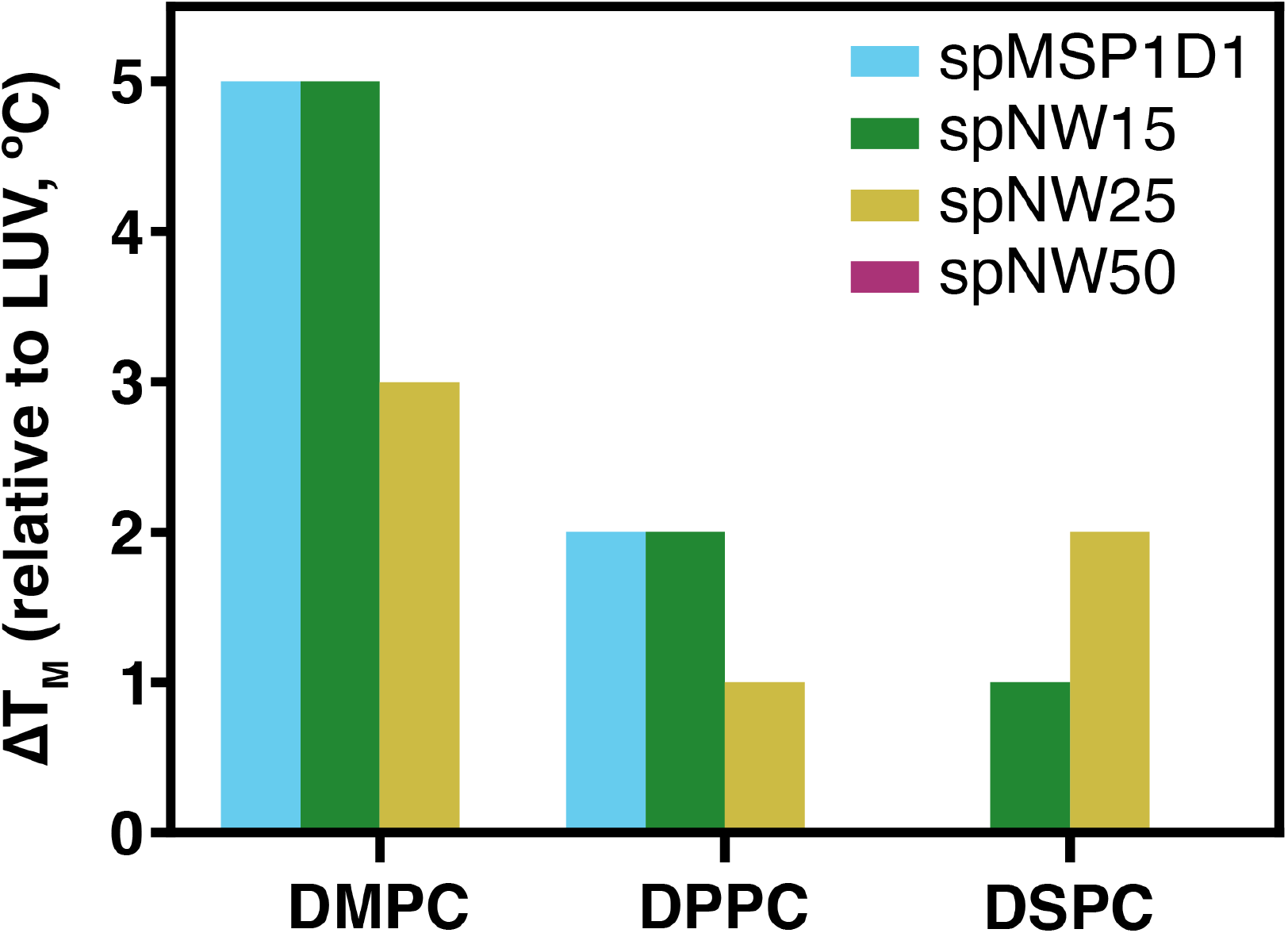
The difference between the main transition (melting) temperature meaured in LUVs and nanodiscs (spMSP1D1, light blue; spNW15, dark green; spNW25, gold; spNW50, magenta) is shown for the saturated lipids examined. Where the colored bar is not visible, the ΔT_*M*_ from the LUV of the same composition is 0°C.

To better understand how lipid unsaturation affects the properties of nanodisc-incorporated lipids, the temperature-dependent ordering of POPC (16:0,18:1) was examined (Fig. 3, Supplemental Fig. S4). As the transition temperature of POPC is -2°C, the transition temperature and cooperativity for this lipid type could not be assayed using the instrument available. Nanodiscs with POPC up to 25 nm showed more order at low temperatures (*i.e*., up to ∼ 20°C) than either spNW50 nanodiscs or LUVs. At higher temperatures, POPC order diverges based on nanodisc size with less ordering in larger nanodiscs; POPC order in spNW50 matches LUVs throughout the temperature spectrum (Fig. 3), as seen with DPPC.

### 3.2. A lipid extract demonstrates complex behavior with regards to nanodisc size

Cell membranes are composed of many lipid types, which are known to allosterically modify membrane proteins. Therefore, structural and biophysical studies involving nanodiscs have frequently used complex lipid mixtures such as soy polar lipids. Soy polar lipids is a lipid extract that contains zwitterionic and anionic headgroups as well as saturated and unsaturated acyl chains [48]. This lipid mixture has been shown to support the function of many membrane proteins, and is therefore frequently used for structural studies involving nanodiscs [40–45]. The properties of these lipids in nanodiscs, however, has rarely been studied to date.

Nanodiscs filled with soy polar lipids show a different trend compared to the single lipid nanodiscs described above. Compared to LUVs, all nanodisc sizes show more ordered lipids across the temperature spectrum examined (4-40°C), including spNW50 (Fig. 6, Supplemental Fig. S5). At the lowest temperature, there is a trend towards larger nanodiscs being less ordered than smaller nanodiscs, which is the opposite of what is observed for single lipid nanodiscs (Fig. 3). As temperature increases, there is a divergence with spNW25. At higher temperatures, lipid order is higher in spNW25 compared to spMSP1D1 and spNW15, and lower in spNW50. Even so, the LUVs containing soy polar extract lipids shows drastically lower order than any nanodisc size studied here. Overall, the behavior of this complex lipid mixture is strikingly different than the trends observed for single lipids discussed above.

**Figure 6:**
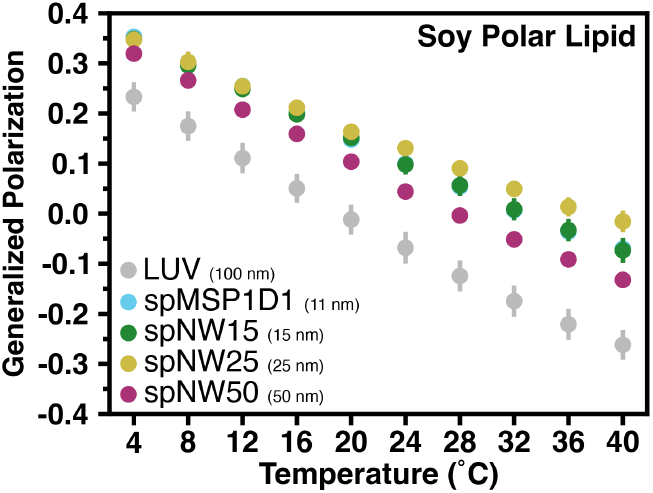
Generalized polarization (GP) values as a function of temperature for soy polar lipids. Each data point represents the average of three independent measurements with the error bars representing the standard deviation of the the three measurements. Data for nanodiscs are colored as light blue for spMSP1D1 (11 nm), dark green for spNW15 (15 nm), gold for spNW25 (25 nm), and magenta for spNW50 (50 nm). Data for LUVs are colored gray. Where data for spMSP1D1 (light blue) is not visible in this plot, data from spNW15 (dark green) overlie due to similarity in the GP values and size of data points.

## 4. Discussion

This study examines the properties of lipids incorporated into nanodiscs of different size to test the long-standing hypothesis that the lipid bilayer in large nanodiscs should behave the same as an unperturbed lipid bilayer [19]. This was achieved by using a circularized nanodisc technology that enables formation of very large nanodiscs. The results demonstrate that as nanodisc size increases, the properties of incorporated lipids show better agreement with LUVs and, in the case of DPPC, DSPC, and POPC, lipids incorporated into spNW50 (50 nm) nanodiscs, show complete agreement with LUVs.

While this data supports the hypothesis that larger nanodiscs support a larger unperturbed lipid bilayer area, the initial model proposed by Sligar and colleagues [19] maybe be incomplete. The model assumed that the volume of perturbed lipids was fixed and ended approximately 1.5 nm from the nanodisc rim. If the perturbed volume was fixed, then the nanodiscs studied here would have approximately 77, 83, 90, and 95% unperturbed lipids (in spMSP1D1, spNW15, spNW25, and spNW50, respectively). Given this, the expected deviation in nanodisc properties compared to LUVs should decrease monotonically, but this is not observed. In fact, in every case examined here, the lipid packing and transition temperatures for spMSP1D1 and spNW15 are indistinguishable. Even in the case of spNW25 and spNW50, for the saturated lipid types outside of transition points, the lipid packing is virtually indistinguishable. Recently, it has been proposed that lipids in a nanodisc do not exist as wholly “perturbed” or “unperturbed”, but rather on a spectrum of perturbation dependent on the distance of the lipid from the nanodisc rim [26]. This idea is supported by multiple simulation studies of MSP-based nanodiscs [21–23] and could explain why there is not a linear trend towards agreement with LUV properties as nanodisc size increases.

Interestingly, there is a dichotomy in the results between single lipid nanodiscs and nanodiscs filled with a complex lipid mixture. While nanodiscs filled with a single lipid follow a simple trend (*i.e*., larger nanodiscs are more similar to LUVs, spNW50 is equivalent to LUVs), this is not observed for soy polar extract nanodiscs. The lipid packing in even spNW50 is drastically different from that observed in LUVs (Fig. 6). Possible reasons for this difference include: (1) preferential incorporation of certain lipid types into the nanodisc via interaction with the MSP scaffold or the SpyCatcher-SpyTag construct, (2) asymmetric distribution of lipids within the nanodisc or LUVs, or (3) asymmetric co-distribution of Laurdan and lipids within the nanodisc. Recently, it was demonstrated that temperature [49] as well as lipid geometry [50] can alter the composition of lipids incorporated into nanodiscs, suggesting that certain lipids preferentially interact with the MSP scaffold or, perhaps, lipids with certain geometries are better accommodated by the curvature induced by the nanodisc scaffold. Therefore, it is plausible that the lipids present in nanodiscs made from soy polar lipid are not the same as those present in LUVs. Indeed, in a recent study using an ER-like membrane, it was shown, via quantitative mass spectromety, that nanodiscs become enriched in cholesterol and depleted in sphingomyelin compared to the native composition [50]. Therefore, nanodiscs affect both the lipid composition as well as the fluid properties of lipid bilayers, which has far reaching implications in structural biology and biophysics, especially in studies where modulatory or necessary lipid species are assumed to be present based on the input composition. It is also entirely possible that specific interactions between lipids and an incorporated transmembrane membrane protein will lead to enrichment of certain lipids in the nanodisc, further altering the composition of lipids within the nanodisc itself. Therefore, the lipid environment of membrane proteins reconstituted in nanodiscs with complex lipid mixtures is likely different from liposomes, even when using large nanodiscs.

It remains unclear whether lipid mixtures asymmetrically distribute within nanodiscs. Simulation studies [21–23, 26] have repeatedly shown spatially-dependent changes in thickness and curvature in nanodiscs, using both all-atom and coarse-grained approaches. Certain lipids could interact favorably with the thin and highly curved edge of the nanodisc rim while other lipids could preferably insert into the thick nanodisc interior. In addition, the MSP contains several charged residues which could drive both favorable and unfavorable interactions between lipids and the nanodisc rim. Certainly, more work is needed to understand what underpins specific lipid-MSP interactions in order to interpret the biophysical data being generated using nanodiscs.

Laurdan was chosen for this study as it is a well-known solvatochromatic dye with well-characterized properties. Importantly, it has been shown repeatedly to partition equally between ordered and disordered phases across a range of lipids [38, 51, 52]. There is also evidence that the presence of a membrane protein in the lipid volume does not alter Laurdan spectra [53]. The current study assumes that Laurdan distributes evenly across the nanodisc lipid bilayer and is reporting on lipid properties in all areas. Nanodiscs, however, are spatially heterogeneous [21–23, 26] with varying thickness and curvature. It is not apparent *a priori* whether Laurdan would distribute evenly across the surface of the nanodisc and, indeed, this question has been raised previously [29]. It is encouraging that similar results were obtained when using different solvatochromatic dyes with different physical properties [29], but further work is needed to understand how Laurdan behaves in the nanodisc environment. Molecular simulations of di-4-ANEPPDHQ, a solvatochromatic, lipophilic fluorophore similar to Laurdan, showed that there can be spatial preference of this positively charged dye for the nanodisc rim in the presence of anionic phospholipids [54]. The same heterogenous distribution was not observed in POPC-only nanodiscs, such as those studied here.

A natural question that follows from these results, due to the central role of nanodiscs in the recent boom of cryo-EM structures is: “What is the optimal nanodisc size for membrane protein structures?”. The equilibrium structure of the ligand-gated ion channel, ELIC, is altered in different size nanodiscs [55], through either altered lipid properties or direct interaction with the nanodisc scaffold. The greatest agonist-dependent changes in ELIC were observed in the largest nanodisc tested, a 15 nm circularized nanodisc (spNW15). Similarly, structures of the ion channel, TRPV3, in 11 nm [56] or 30 nm [57] circularized nanodiscs captured different conformations compared to smaller non-circularized nanodiscs, providing insights into the mechanisms of heat activation and opening of this ion channel. These studies suggest that nanodisc size can impact the structure of membrane proteins, and it remains to be determined if the properties of the lipid bilayer are contributing factors. However, there is also evidence that membrane proteins preferentially interact with the nanodisc rim and are stabilized by interactions with the scaffold [58, 59]. Therefore, it is unclear what the “optimal” nanodisc size is for membrane protein structure determination and it is very likely that it is dependent on the membrane protein of interest and its interaction with the nanodisc itself.

## 5. Conclusion

Nanodiscs remain a valuable tool for biophysical and structural studies of membrane proteins. It has long been known, however, that lipids incorporated into nanodiscs have altered physical properties compared to unperturbed lipid bilayers [19, 29]. Using circularized nanodiscs that encompass a large range of sizes, this study demonstrates that as nanodisc size increases, the physical properties of the lipid bilayer such as lipid packing, melting temperature, and cooperativity better approximate that of LUVs for single lipid compositions. With complex lipid mixtures, such as soy polar lipids, this trend does not hold and spNW50 does not recover the physical properties of an LUV. This could be due to preferential enrichment of certain lipid species within the nanodisc or asymmetric distribution of lipids within the nanodisc, but further work is needed to test this. Overall, these results indicate that larger nanodiscs support a more unperturbed membrane environment, but the extent to which this is true will depend on the types and combinations of lipids used.

## Author Contributions

M.J.A. and W.W.C designed the research. M.J.A. and V.D. expressed and purified the circularized nanodiscs. M.J.A. and D.F. prepared the lipid-filled nanodiscs and liposomes as well as performed and analyzed the fluorometric assay. M.J.A. wrote the analysis code and analyzed the results of the fluorometric assay. M.J.A., V.D., D.F., and W.W.C. wrote and edited the manuscript.

## Acknowledgments

The authors wish to thank Dr. Alex Evers for the generous use of his spectrophotomer for these studies. The authors also wish to thank the National Institutes of Health (K08-GM152844 to M.J.A. and R35-GM137597 to W.W.C.) and the Foundation for Anesthesia Education and Research (Mentored Research Training Grant to M.J.A.) for the generous support which enabled these studies.

## Data Availability

A static version of the Python-based data analysis script, together with the raw data used in this study, is deposited at DRYAD. As mentioned in 2.5, a live version of the data analysis script is located at: https://github.com/mjarcario/Laurdan-Data-Extraction/tree/main.

## References

[1] I. Levental, E. Lyman, Regulahtion of membrane protein structure and function by their lipid nano-environment, Nat. Rev. Mol. Cell Biol. 24 (2023) 107–122.

[2] N. Thakur, A. P. Ray, L. Sharp, B. Jin, A. Duong, N. G. Pour, S. Obeng, A. V. Wijesekara, Z.-G. Gao, C. R. McCurdy, K. A. Jacobson, E. Lyman, M. T. Eddy, Anionic phospholipids control mechanisms of GPCR-G protein recognition, Nat. Commun. 14 (2023) 794.

[3] N. Thakur, A. P. Ray, B. Jin, N. P. Afsharian, E. Lyman, Z.-G. Gao, K. A. Jacobson, M. T. Eddy, Membrane mimetic-dependence of GPCR energy landscapes, Structure 32 (2024) 1–13.

[4] T. H. Bayburt, Y. V. Grinkova, S. G. Sligar, Self-assembly of discoidal phospholipid bilayer nanoparticles with membrane scaffolding proteins, Nano Lett. 2 (2002) 853–856.

[5] J. Frauenfeld, R. Löving, J.-P. Armache, A. F.-P. Sonnen, F. Guettou, P. Moberg, C. Jegerschöld, A. Flayhan, J. A. G. Briggs, H. Garoff, C. Löw, Y. Cheng, P. Nordlund, A saposin-lipoprotein nanoparticle system for membrane proteins, Nat. Methods 13 (2016) 345–351.

[6] M. L. Carlson, J. W. Young, Z. Zhao, L. Fabre, D. Jun, J. Li, H. S. Dhupar, I. Wason, A. T. Mills, J. T. Beatty, J. S. Klassen, I. Rouiller, F. Duong, The Peptidisc, a simple method for stabilzing membrane proteins in detergent-free solution, eLife 7 (2018) e34085.

[7] Z. Zhao, M. Zhang, J. M. Hogle, W. M. Shih, G. Wagner, M. L. Nasr, DNA-corralled nanodiscs for the structural and functional characterization of membrane proteins and viral entry, J. Am. Chem. Soc. 140 (2018) 10639–10643.

[8] T. Ravula, N. Z. Hardin, A. Ramamoorthy, Polymer nanodiscs: Advantages and limitations, Chem. Phys. Lipids 219 (2019) 45–49.

[9] M. Overduin, M. Esmaili, Memtein: The fundamental unit of membrane-protein structure and function, Chem. Phys. Lipids 218 (2019) 73–84.

[10] M. D. Farrelly, L. L. Martin, S. H. Thang, Polymer nanodiscs and their bioanalytical potential, Chem. Eur. J. 27 (2021) 12922–12939.

[11] I. G. Denisov, S. G. Sligar, Nanodiscs for structural and functional studies of membrane proteins, Nat. Struct. Mol. Biol. 23 (2016) 481–486.

[12] R. G. Efremov, C. Gatsogiannis, S. Raunser, Lipid nanodiscs as a tool for high-resolution structure determination by single-particle cryo-EM, Methods Enzymol. 594 (2017) 1–30.

[13] U. Günsel, F. Hagn, Lipid nanodiscs for high-resolution NMR studies of membrane proteins, Chem. Rev. 122 (2022) 9395–9421.

[14] M. A. McLean, I. G. Denisov, Y. V. Grinkova, S. G. Sligar, Dark, ultra-dark, and ultra-bright nanodiscs for membrane protein investigations, Anal. Biochem. 607 (2020) 113860.

[15] A. Nath, P. K. Koo, E. Rhoades, W. M. Atkins, Allosteric effects on substrate dissociation from cytochrome P450 3A4 in nanodiscs observed by ensemble and single-molecule fluorescence spectroscopy, J. Am. Chem. Soc. 130 (2008) 15746–15747.

[16] F. Nakagawa, M. Kikkawa, S. Chen, Y. Miyashita, N. Hamaguchi-Suzuki, M. Shibuya, S. Yamashita, L. Nagase, S. Yasuda, M. Shiroshi, T. Senda, K. Ito, T. Murata, S. Ogasawara, Anti-nanodisc antibodies specifically capture nanodiscs and facilitate molecular interaction kinetics studies for membrane protein, Sci. Rep. 13 (2023) 11627.

[17] X. Ye, M. A. McLean, S. G. Sligar, Conformational equilibrium of talin is regulated by anionic lipids, BBA Biomembr. 1858 (2016) 1833–1840.

[18] Q. Ren, S. Zhang, H. Bao, Circularized fluorescent nanodiscs for probing protein-lipid interactions, Commun. Biol. 5 (2022) 507.

[19] I. G. Denisov, M. A. McLean, A. W. Shaw, Y. V. Grinkova, S. G. Sligar, Thermotropic phase transition in soluble nanoscale lipid bilayers, J. Phys. Chem. B 109 (2005) 15580–15588.

[20] D. Martinez, M. Decossas, J. Kowal, L. Frey, H. Stahlberg, E. J. Dufourc, R. Riek, B. Habenstein, S. Bibow, A. Loquet, Lipid internal dynamics probed in nanodiscs, ChemPhysChem 18 (2017) 2651–2657.

[21] Y. Qi, J. Lee, J. B. Klauda, W. Im, CHARMM-GUI Nanodisc Builder for modeling and simulation of various nanodisc systems, J. Comput. Chem. 40 (2019) 893–899.

[22] T. Bengtsen, V. L. Holm, L. R. Kjølbye, S. R. Midtgaard, N. T. Johansen, G. Tesei, S. Bottaro, B. Schiøtt, L. Arleth, K. Lindorff-Larsen, Structure and dynamics of a nanodisc by integrating NMR, SAXS and SANS experiments with molecular dynamics simulations, eLife 9 (2020) e56518.

[23] I. Schachter, C. Allolio, G. Khelashvili, D. Harries, Confinement in nanodiscs anisotropically modifies lipid bilayer elastic properties, J. Phys. Chem. B 124 (2020) 7166–7175.

[24] L. R. Kjølbye, L. De Maria, T. A. Wassenaar, H. Abdizadeh, S. J. Marrink, J. Ferkinghoff-Borg, B. Schiøtt, General protocol for constructing molecular models of nanodiscs, J. Chem. Inf. Model. 61 (2021) 2869– 2883.

[25] A. Debnath, L. V. Schäfer, Structure and dynamics of phospholipid nanodiscs from all-atom and coarse-grained simulations, J. Phys. Chem. B 119 (2015) 6991–7002.

[26] P. Stepien, B. Augustyn, C. Poojari, W. Galan, A. Polit, I. Vattulainen, A. Wisnieska-Becker, T. Rog, Complexity of seemingly simple nanodiscs, BBA Biomembr. 1862 (2020) 183420.

[27] P.-S. Ho, T.-Y. Kao, C.-C. Li, Y.-J. Lan, Y.-C. Lai, Y.-W. Chiang, Nanodisc lipids exhibit singular behaviors impying critical phenomena, Langmuir 38 (2022) 15372–15383.

[28] A. P. Bali, I. D. Sahu, A. F. Craig, E. E. Clark, K. M. Burridge, M. T. Dolan, C. Dabney-Smith, D. Konkolewicz, G. A. Lorigan, Structural characterization of styrene-maleic acid copolymer-lipid nanoparticles (SMALPs) using EPR spectroscopy, Chem. Phys. Lipids (2019) 6–13.

[29] L. M. Real Hernandex, I. Levental, Lipid packing is disrupted in copolymeric nanodiscs compared with intact membranes, Biophys. J. 122 (2023) 2256–2266.

[30] Y. V. Grinkova, I. G. Denisov, S. G. Sligar, Engineering extended membrane scaffold proteins for self-assembly of soluble nanoscale lipid bilayers, Protein Eng. Des. Sel. 23 (2010) 843–848.

[31] M. L. Nasr, D. Baptista, M. Strauss, Z. J. Sun, S. Grigoriu, S. Huser, A. Plückthun, F. Hagn, T. Walz, J. M. Hogle, G. Wagner, Covalently circularized nanodiscs for studying membrane proteins and viral entry, Nat. Methods 14 (2017) 49–52.

[32] J. Miehling, D. Goricanec, F. Hagn, A split-intein-based method for the efficient production of circularized nanodiscs for structural studies of membrane proteins, ChemBioChem 19 (2018) 1927–1933.

[33] S. Zhang, Q. Ren, S. J. Novick, T. S. Strutzenberg, P. R. Griffin, H. Bao, One-step construction of circularized nanodiscs using SpyCatcher-SpyTag, Nat. Commun. 12 (2021) 5451.

[34] T. Parasassi, E. Krasnowska, L. Bagatolli, E. Gratton, Laurdan and Prodan as polarity-sensitive fluorescent membrane probes, J. Fluoresc. 8 (1998) 365–373.

[35] A. W. Shaw, M. A. McLean, S. G. Sligar, Phospholipid phase transitions in homogenous nanometer scale bilayer discs, FEBS Lett. 556 (2004) 260–264.

[36] J. J. Dominguez Pardo, J. M. Dörr, M. F. Renne, T. Ould-Braham, M. C. Koorengevel, M. J. van Steenbergen, J. A. Killian, Thremotropic properties of phosphatidylcholine nanodiscs bounded by styrene-maleic acid copolymers, Chem. Phys. Lipids 208 (2017) 58–64.

[37] T. Camp, S. G. Sligar, Nanodisc self-assembly is thermodynamically reversible and controllable, Soft Matter 16 (2020) 5615–5623.

[38] M. Amaro, F. Reina, M. Hof, C. Eggeling, E. Sezgin, Laurdan and di4-ANEPPDHQ probe different properties of the membrane, J. Phys. D Appl. Phys. 50 (2017) 134004.

[39] R. Koynova, M. Caffrey, Phases and phase transitions of the phosphatidylcholines, BBA Biomembr. (1998) 91–145.

[40] N. Sengupta, A. K. Mondal, S. Mishra, K. Chattopadhyay, S. Dutta, Single-particle cryo-EM reveals conformational variability of the oligomeric VCC β-barrel pore in a lipid bilayer, J. Cell. Biol. 220 (2021) e202102035.

[41] U. Goswami, M. M. Rahman, J. Teng, R. E. Hibbs, Structural interplay of anesthetics and paralytics on muscle nicotinic receptors, Nat. Commun. 14 (2023) 3169.

[42] Y. Gao, E. Cao, D. Julius, Y. Cheng, TRPV1 structures in nanodiscs reveal mechanisms of ligand and lipid action, Nature 534 (2016) 347– 351.

[43] A. F. Kintzer, E. M. Green, P. K. Dominik, M. Bridges, J.-P. Armache, D. Deneka, S. S. Kim, W. Hubbell, A. K. Kossiakoff, Y. Cheng, R. M. Stroud, Structural basis for the activation of voltage sensor domains in an ion channel TPC1, Proc. Natl. Acad. Sci. USA 115 (2018) E9095– E9104.

[44] V. Kalienkova, V. C. Mosina, L. Bryner, G. T. Oostergetel, R. Dutzler, C. Paulino, Stepwise activation mechanism of the scramblase nhT-MEM16 revealed by cryo-EM, eLife 8 (2019) e44364.

[45] A. Kumar, S. Basak, S. Rao, Y. Gicheru, M. L. Mayer, M. S. P. Sansom, S. Chakrapani, Mecahnisms of activation and desensitization of full-length glycine receptor in lipid nanodiscs, Nat. Commun. 11 (2020) 3752.

[46] J. J. Dominguez Pardo, J. M. Dörr, M. F. Renne, T. Ould-Braham, M. C. Koorengevel, M. J. van Steenbergen, J. A. Killian, Thermotropic properties of phosphatidylcholine nanodiscs bounded by styrene-maleic acid copolymers, Chem. Phys. Lipids 208 (2017) 58–64.

[47] N. Färber, C. Westerhausen, Broad lipid phase transitions in mammalian cell membranes measured by Laurdan fluorescence spectroscopy, BBA Biomembr. (2022) 183794.

[48] T. Wang, E. G. Hammond, W. R. Fehr, Phospholipid fatty acid composition and stereospecific distribution of soybeans with a wide range of fatty acid composition, J. Am. Oil Chem.’ Soc. 74 (1997) 1587–1594.

[49] M. T. Odenkirk, G. Zhang, M. T. Marty, Do nanodisc assembly conditions affect natural lipid uptake?, J. Am. Soc. Mass Spectrom. 34 (2023) 2006–2015.

[50] M. C. Sarcinella, J. D. Jones, M. J. Sorensen, S. A. Edgcombe, B. T. Ruotolo, R. T. Kennedy, R. C. Bailey, Lipid curvature and fluidity influence lipid incorporation disparities in nanodiscs, Anal. Chem. 97 (2025) 2883–2889.

[51] T. Parasassi, G. De Stasio, A. d’Ubaldo, E. Gratton, Phase fluctuation in phospholipid membranes revealed by Laurdan fluorescence, Biophys. J. 57 (1990) 1179–1186.

[52] H.-J. Kaiser, D. Lingwood, I. Levental, J. L. Sampaio, L. Kalvodova, L. Rajendran, K. Simons, Order of lipid phases in model and plasma membranes, Proc. Natl. Acad. Sci. USA 106 (2009) 16645–16650.

[53] J. Dinic, H. Biverståhl, L. Mäler, I. Parmryd, Laurdan and di-4-ANEPPDHQ do not respond to membrane-inserted peptides and are good probes for lipid packing, BBA Biomembr. 1808 (2011) 298–306.

[54] A. Chmielińska, P. Stepien, P. Bonarek, M. Girych, G. Enkavi, T. Rog, M. Dziedzicka-Wasylewska, A. Polit, Can di-4-ANEPPDHQ reveal the structural differences between nanodiscs and liposomes?, BBA Biomembr. 1863 (2021) 183649.

[55] V. Dalal, M. J. Arcario, J. T. Petroff II, B. K. Tan, N. M. Dietzen, M. J. Rau, J. A. J. Fitzpatrick, G. Brannigan, W. W. L. Cheng, Lipid nanodisc scaffold and size alter the structure of a pentameric ligand-gated ion channel, Nat. Commun. 15 (2024) 25.

[56] K. D. Nadezhdin, A. Neuberger, Y. A. Trofimov, N. A. Krylov, V. Sinica, N. Kupko, V. Vlachova, E. Zakharian, R. G. Efremov, A. I. Sobolevsky, Structural mechanism of heat-induced opening of a temperature-sensitive TRP channel, Nat. Struct. Mol. Biol. 28 (2021) 564–572.

[57] K. D. Nadezhdin, A. Neuberger, L. S. Khosrof, I. A. Talyzina, J. Khau, M. V. Yelshanskaya, A. I. Sobolevsky, TRPV3 activation by different agonists accompanied by lipid dissociation from the vanilloid site, Sci. Adv. 10 (2024) eadn2453.

[58] C. A. López, M. F. Swift, X.-P. Xu, D. Hanein, N. Volkmann, S. Gnanakaran, Biophysical characterization of a nanodisc with and without BAX: An integrative study using molecular dynamics simulations and cryo-EM, Structure 27 (2019) 988–999.

[59] S. Orioli, C. G. H. Hansen, L. Arleth, Ab initio determination of the shape of membrane proteins in a nanodisc, Acta Cryst. D77 (2021) 176– 193.

