## Supplemental Data for "Examining the Thermotropic properties of Large, Circularized Nanodiscs"

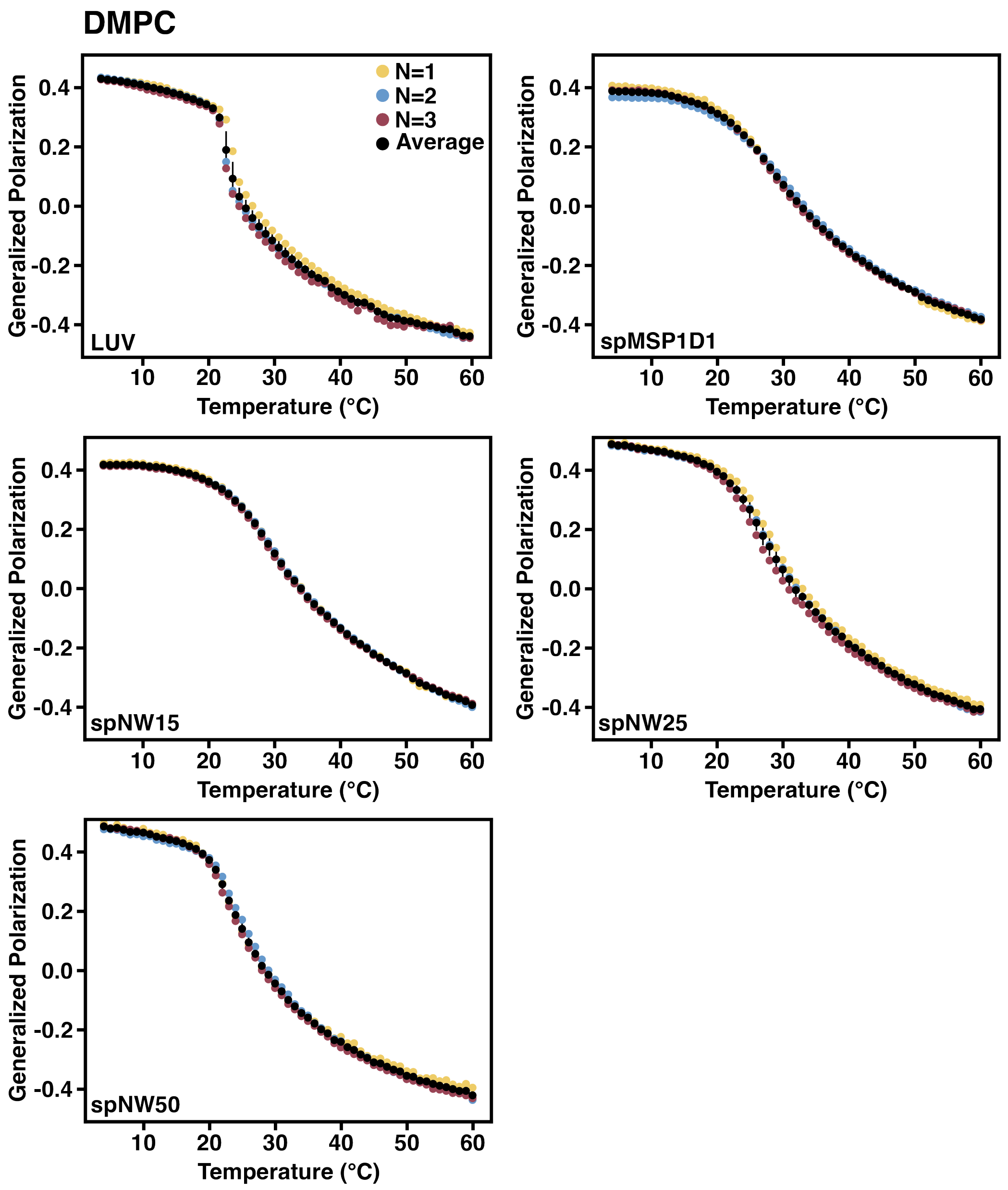


Figure S1: Generalized polarization versus temperature for all three independent DMPC replicates (yellow, blue, and red dots). Average across the three independent replicates is shown as black dots with the error bar representing the standard deviation across the three replicates.

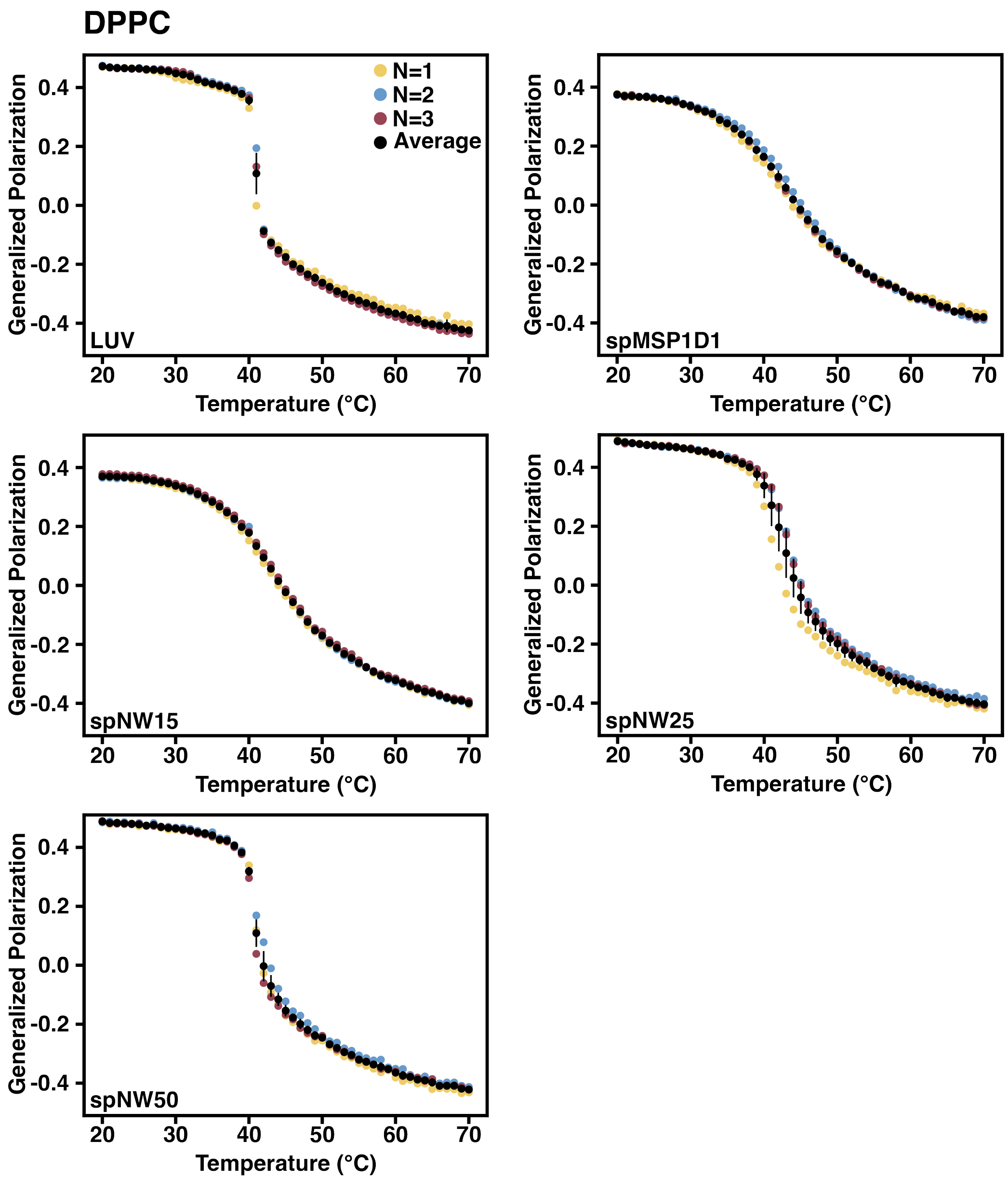


Figure S2: Generalized polarization versus temperature for all three independent DPPC replicates (yellow, blue, and red dots). Average across the three independent replicates is shown as black dots with the error bar representing the standard deviation across the three replicates.

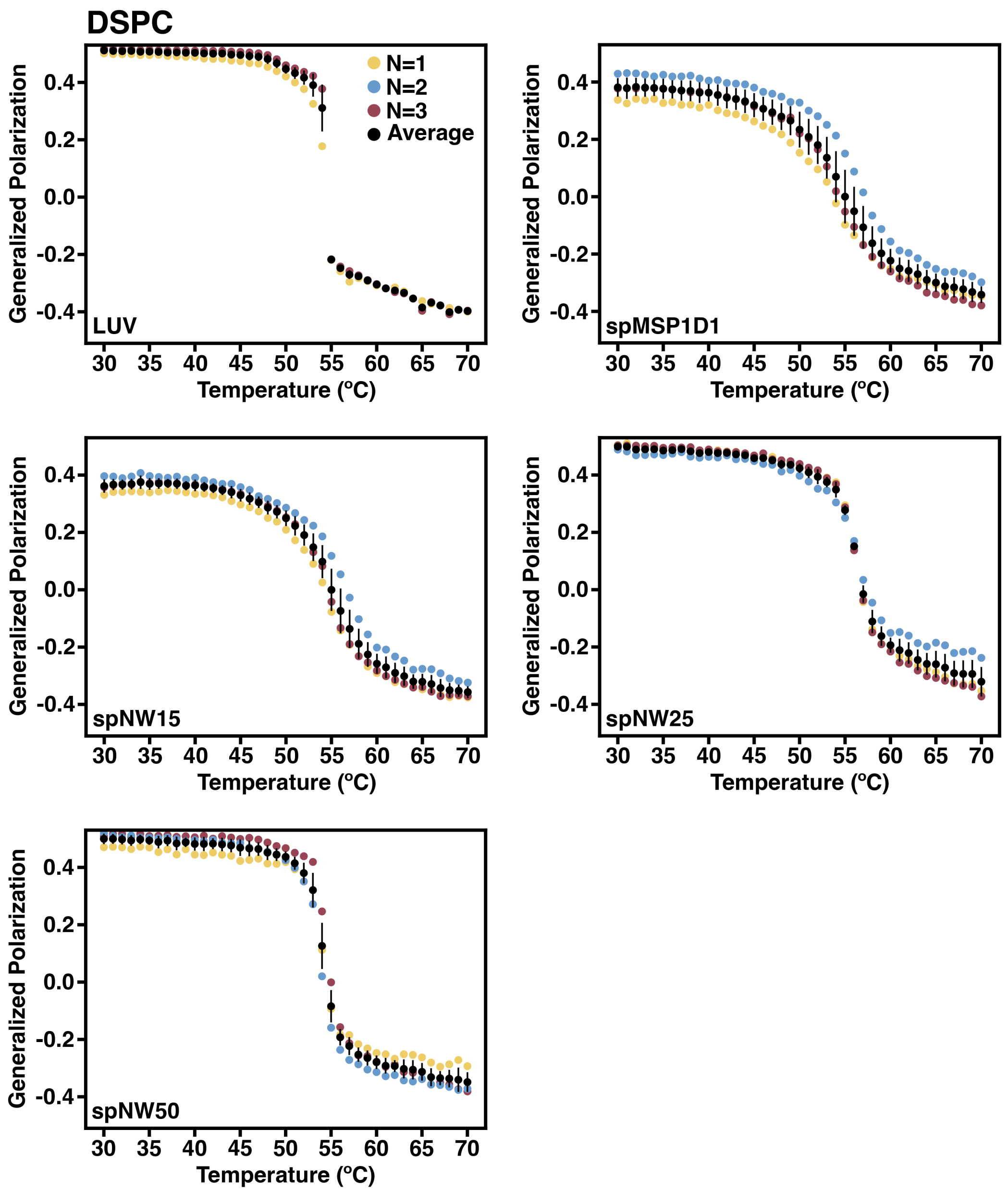


Figure S3: Generalized polarization versus temperature for all three independent DPPC replicates (yellow, blue, and red dots). Average across the three independent replicates is shown as black dots with the error bar representing the standard deviation across the three replicates.

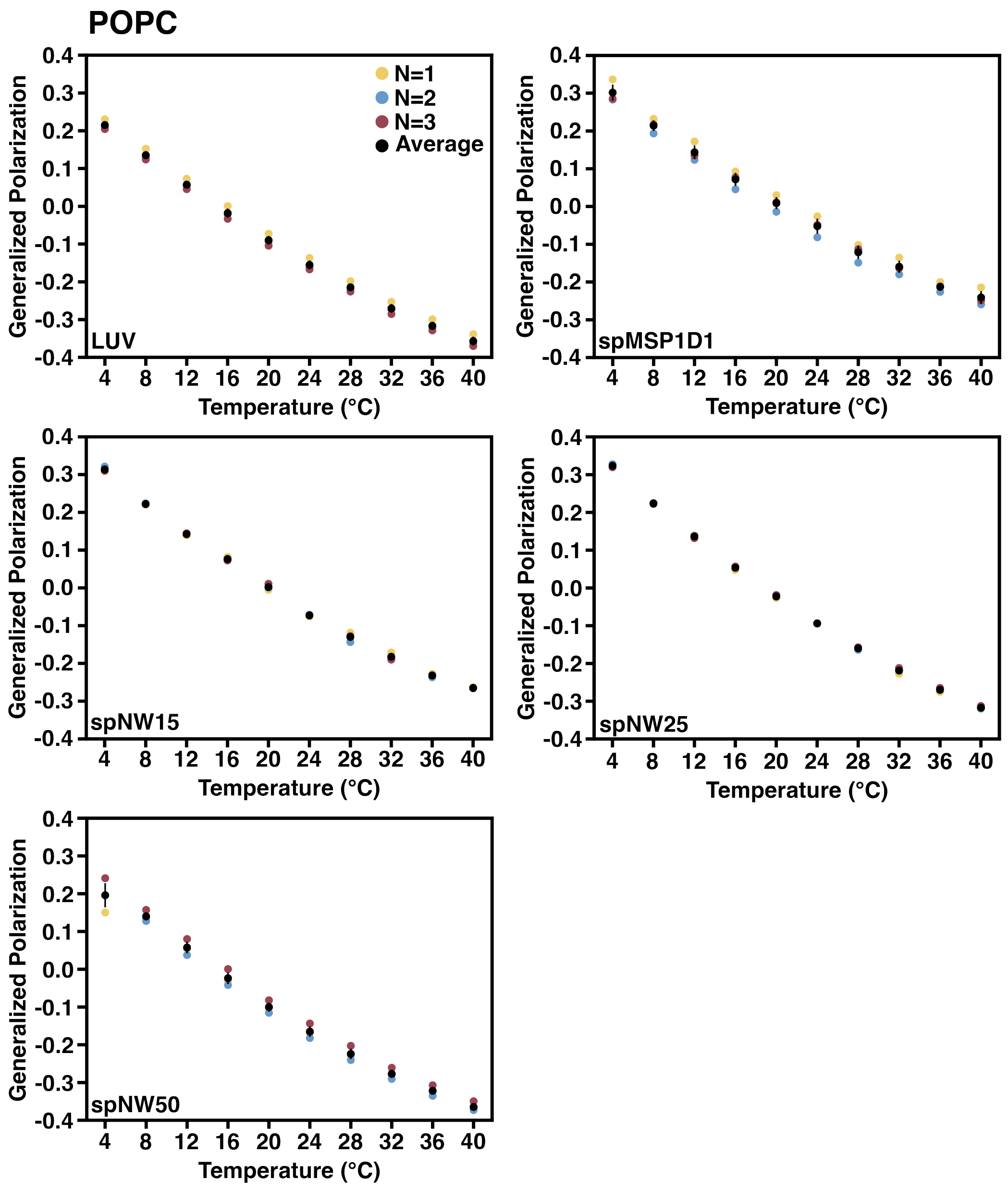


Figure S4: Generalized polarization versus temperature for all three independent POPC replicates (yellow, blue, and red dots). Average across the three independent replicates is shown as black dots with the error bar representing the standard deviation across the three replicates.

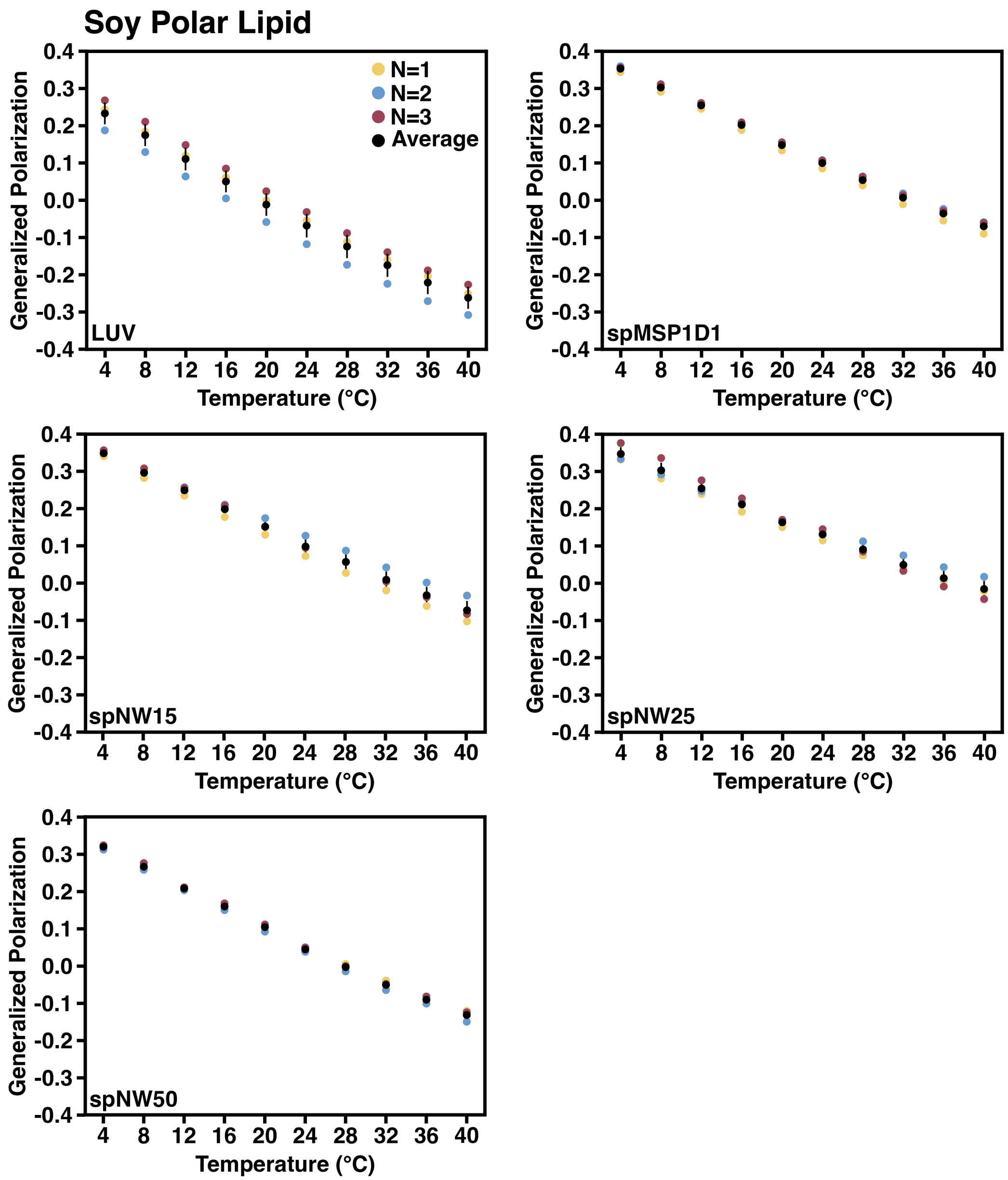


Figure S5: Generalized polarization versus temperature for all three independent soy polar extract replicates (yellow, blue, and red dots). Average across the three independent replicates is shown as black dots with the error bar representing the standard deviation across the three replicates.
